# Complete structure of the core signalling unit of the *E. coli* chemosensory array in an optimised minicell strain

**DOI:** 10.1101/723692

**Authors:** Alister Burt, C. Keith Cassidy, Peter Ames, Maria Bacia-Verloop, Megghane Baulard, Karine Huard, Zaida Luthey-Schulten, Ambroise Desfosses, Phillip J. Stansfeld, William Margolin, John S. Parkinson, Irina Gutsche

## Abstract

Motile bacteria sense chemical gradients with transmembrane receptors organised in supramolecular signalling arrays.^1,2^ Understanding stimulus detection and transmission at the molecular level requires precise structural characterisation of the array building block known as a core signalling unit (CSU). Here we introduce a novel *E. coli* strain that forms small minicells possessing extended and highly ordered chemosensory arrays. We provide a three-dimensional (3D) map of a complete CSU at ~16 Å resolution by cryo-electron tomography (cryo-ET) and subtomogram averaging. This map, combined with previously determined high resolution structures and molecular dynamics simulations, yields an atomistic model of the membrane-bound CSU and enables spatial localisation of its signalling domains. Our work thus offers a solid structural basis for interpretation of existing data and design of new experiments to elucidate signalling mechanisms within the CSU and larger array.

---

Bacteria survive and proliferate by sensing changes in their environment and responding through metabolic adaptation or locomotion. Chemotactic bacteria, for example, monitor attractant and repellent concentration gradients and promote movement towards favorable niches. The chemosensory arrays that mediate this behavior are assembled from CSUs that contain two trimers of receptor dimers (ToDs) interconnected at their cytoplasmic tips by a dimeric histidine autokinase CheA and two copies of a coupling protein CheW, which links CheA activity to receptor control (Fig. 1a, b, c). Chemoeffector binding to the periplasmic domain of the receptors triggers a signalling cascade of intracellular phosphorylation events which ultimately regulate the direction of the cell’s flagellar motors. To allow appropriate sensory responses over a wide range of chemoeffector concentrations, receptor sensitivity is continuously tuned through the reversible methylation of receptors, known as methyl-accepting chemotaxis proteins or MCPs. *E. coli* has four canonical MCPs that share a common functional architecture (Fig. 1a); the two numerically predominant ones are Tar (aspartate and maltose sensor) and Tsr (serine and autoinducer 2 sensor). Changes in ligand occupancy of their periplasmic ligand-binding domains trigger conformational rearrangements that propagate through the inner membrane to the HAMP domain, a signalling element found in most microbial chemoreceptors and sensory kinases.^3^ The HAMP domain couples extracellular input to intracellular output by relaying stimulus information through an extended methylation helix (MH) bundle, via a flexible region and glycine hinge, to the signalling tips where it is further transmitted to CheA and CheW to affect kinase activity. CheA functions as a homodimer with each monomer containing five domains (referred to as P1-P5) connected by flexible linkers (Fig. 1b).

**Figure 1:**
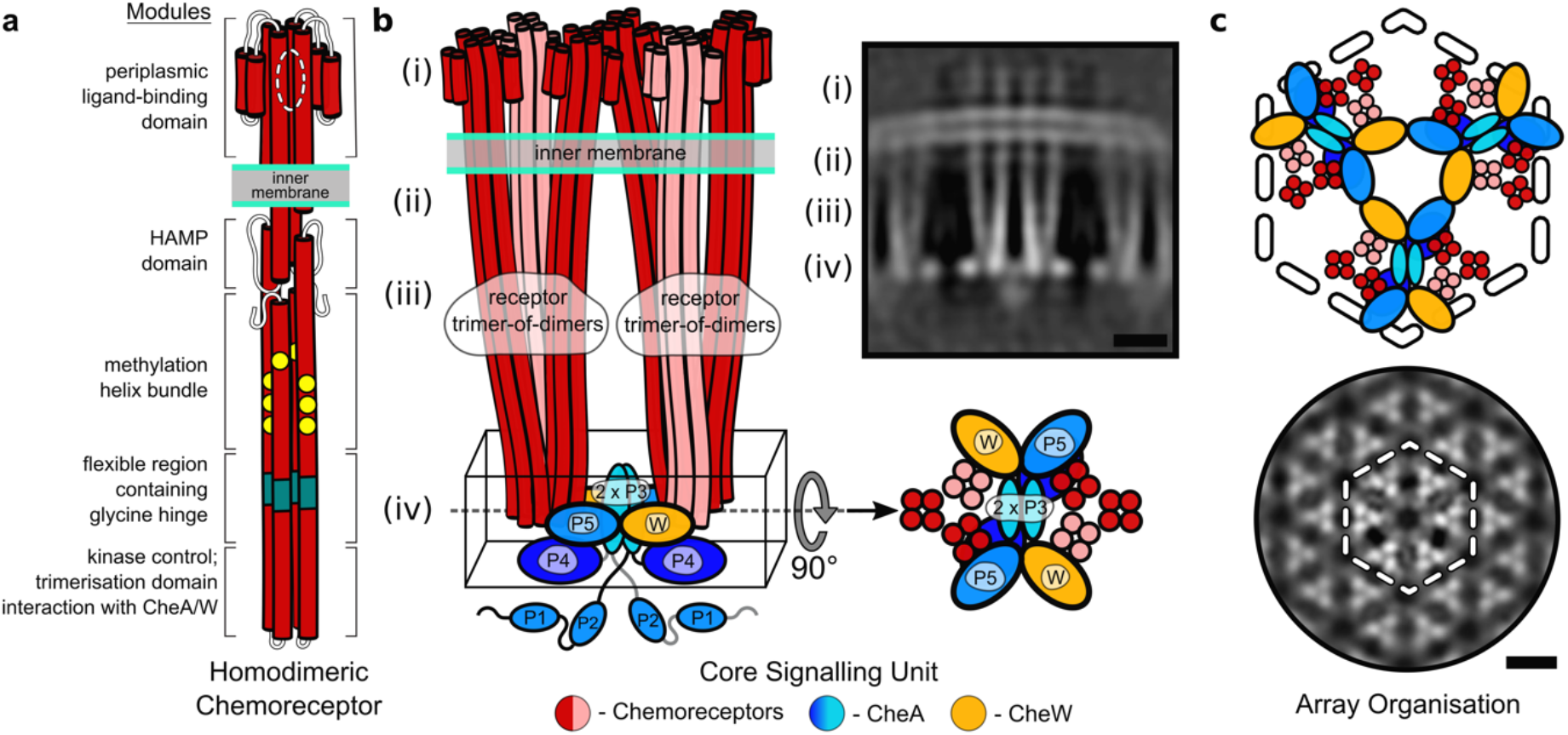
Overview of chemoreceptor, core signalling unit and chemoreceptor array architectures. Schematic representation of homodimeric chemoreceptor structure. Red cylinders represent α-helical secondary structures drawn approximately to scale, flexible hinges are drawn as thin wavy strings, important regions discussed further in the text are highlighted (methylation sites as yellow circles, glycine hinges as teal cylinders). Regions encompassed in square brackets are the periplasmic ligand-binding domain (PP), the HAMP domain (HAMP), the methylation-helix bundles (MH), a flexible region containing the glycine hinge (GH) and the trimerisation and kinase control domain (KC) which is the site of interaction with CheA and CheW. (b – left) Two receptor ToDs interact with CheA and CheW to form a CSU shown from the side. Two MCP dimers in the CSU are shown in salmon for perspective. CheA is shown in shades of blue, and CheW in gold. CheA.P3, CheA.P4 and CheA.P5 are labelled and have known positions. Positions of CheA.P1 and CheA.P2 are not certain. The baseplate region is boxed. (b – top right) Same as in (b – left) shown as a 5 nm thick projection through the density of our reconstruction. Roman numerals in (b – left) and (b – top right) refer to important regions of the structure – (i) the periplasmic domain, (ii) the HAMP domain, (iii) the methylation-helix bundle and (iv) the baseplate region. (c – top) CheA and CheW from three neighbouring CSUs interact to form chemoreceptor arrays, shown schematically and (c – bottom) as a 5 nm projection through the density of our reconstruction. The same region in (c – top) and (c – bottom) is delimited by a dashed hexagon. In (b – top right) and (c – bottom), protein density is shown in white and scale bars are 10 nm.

The mechanistic details of sensory signal transmission within the CSU remain mysterious, in part because there is as yet no experimentally determined structure, even at low resolution, for a complete CSU. On one hand, cryo-ET structures of complexes composed of receptor cytoplasmic domains, CheA and CheW that are assembled on lipid monolayers provide insights into CSU structure,^4^ but lack the ligand-binding, transmembrane (TM), and HAMP portions of the receptor. On the other hand, cryo-ET maps of intact receptor arrays *in situ* thus far lack this level of detail and become diffuse at membrane-proximal regions, purportedly due to intrinsic flexibility of the signalling domains.^5–8^

While cryo-ET offers the best prospects for understanding the structures of receptors and their signalling arrays,^9–11^ most bacteria are not ideal candidates for direct cryo-ET imaging because their thickness exceeds the practical limit of ~500 nm. In principle this limit can be circumvented by vitreous sectioning^12,13^ or focused ion beam milling^14–16^; however, these methods are expensive, technically demanding, and introduce the additional challenge of identifying sections containing the areas of interest. Bacterial minicells offer a valuable solution to the thickness problem. Minicell-producing strains frequently divide close to one pole of a rod-shaped bacterium to generate small achromosomal cells capable of normal metabolic functions. Such minicells are thus particularly useful for studies of polar regions of the bacterium, precisely where *E. coli* chemoreceptors organize into arrays.^17^

Here we present the WM4196 minicell-producing *E. coli* strain from which we isolated small, healthy-looking minicells suitable for high-resolution analysis of internal structures by cryo-ET and subtomogram averaging. Typical minicell preparations reveal numerous defects, including stripped-off membranes, vesicles, spheroplasts, several concentric membrane layers, and a markedly swollen periplasm.^5,6,18,19^ The WM4196 strain produced a higher proportion of uniform and round minicells, many of which were less than 0.4 µm in diameter (Fig. 2a). In addition, cryo-ET reconstructions revealed flattening of these cells in vitreous ice, presumably caused by the plunge freezing process. This flattening reflects the relative plasticity of the cells and makes the sample thinner and, therefore, more suitable for high-resolution cryo-ET analysis (Fig. 2b). Moreover, we discovered that about 50% of WM4196 minicell cryo-ET reconstructions featured extensive chemosensory arrays (Fig. 2c), in contrast to a previous minicell study in which less than 20% contained visible arrays.^6^ Therefore we decided to characterize the WM4196 strain genetically and biochemically as a potentially useful tool for the structural investigation of chemosensory arrays.

**Figure 2:**
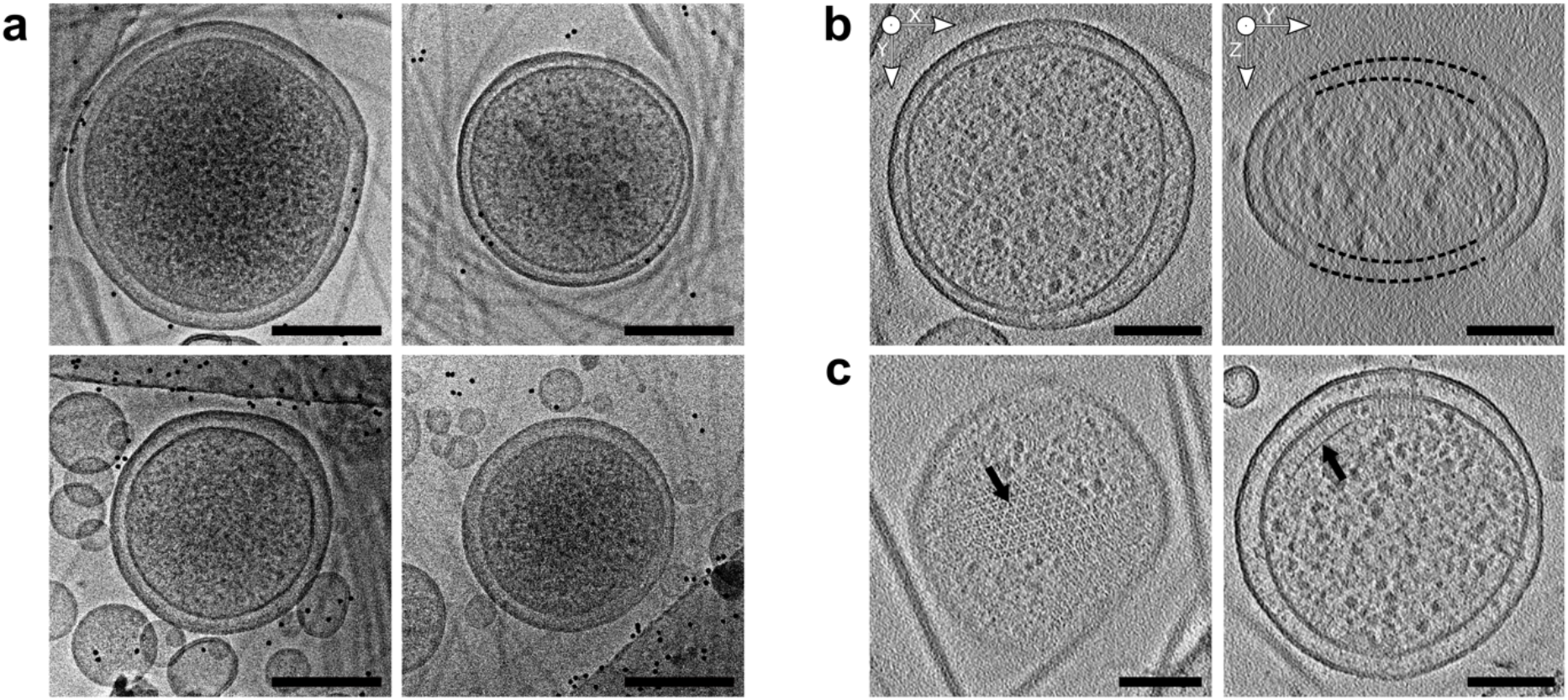
WM4196 minicells are suitable tools for high resolution analysis of chemoreceptor arrays. (a) Representative electron micrographs showing healthy looking WM4196 minicells. (b – left) XY slice and (b – right) YZ slices through tomograms of WM4196 minicells showing that they appear flattened in vitreous ice, yielding thin samples suitable for cryo-ET. Black dotted lines are used to show membrane positions which are not clear in the images due to missing-wedge effects from tomographic reconstruction. (c) slices through tomograms of WM4196 minicells exhibiting presence of chemoeceptor arrays aligned both perpendicular (left) and parallel (right) to the electron beam. Scale bar is 100 nm in all panels.

Strain WM4196 is a distant derivative of a minicell mutant first described over 50 years ago,^20^ into which we introduced the *mreB-A125V* allele (see Methods).^21,22^ Sequence analysis of the *minCDE* region in WM4196 revealed a *minD-G262D* mutation, which in combination with the *mreB* mutation should produce a skinny minicell phenotype.^23^ In addition, sequence comparisons with RP437, a wild-type reference strain for *E. coli* chemotaxis studies,^24^ identified a single base-pair mutation [*pflhDC(−10)g5a*] in WM4196 at the −10 region of the promoter for the *flhDC* genes, whose products promote expression of all flagellar and chemotaxis genes.^25^ To ascertain the contributions of the *mreB*, *minD*, and *pflhDC* alleles to the favorable cryo-ET imaging properties of WM4196 minicells, we constructed RP437 derivatives with combinations of the WM4196 alleles.

First, to examine whether WM4196 minicells were smaller than those made by an RP437 derivative (UU3118) carrying the *mreB-A125V* and *minD-G262D* alleles, we isolated minicells from cultures of WM4196 and UU3118 grown under identical conditions and compared their sizes by analysis of phase contrast light micrographs. WM4196 minicells were overall smaller and had a tighter size distribution than those produced by UU3118 (Supplementary Fig. 1). Although we do not yet know the genetic basis for this size difference, a sequence comparison of the entire RP437 and WM4196 genomes is in progress and should identify candidate genes for subsequent study.

Second, cultures of WM4196 and RP437 derivatives were examined for expression of three principal chemosensory array components: the Tsr and Tar chemoreceptors and the CheA kinase. Levels of all three components were 2- to 3-fold higher in WM4196 than in RP437 (Fig. 3a). Most of the expression increase likely derives from the *flhDC* promoter mutation, but the *mreB and minD* allele combination of WM4196 also seems to contribute to the difference (Fig. 3a). An RP437 derivative (UU3120) carrying the *mreB*, *minD*, and *pflhDC* alleles of WM4196 produced comparably elevated expression of these array components, although the Tar:Tsr ratio was slightly different between the two strains (Fig. 3a).

**Figure 3:**
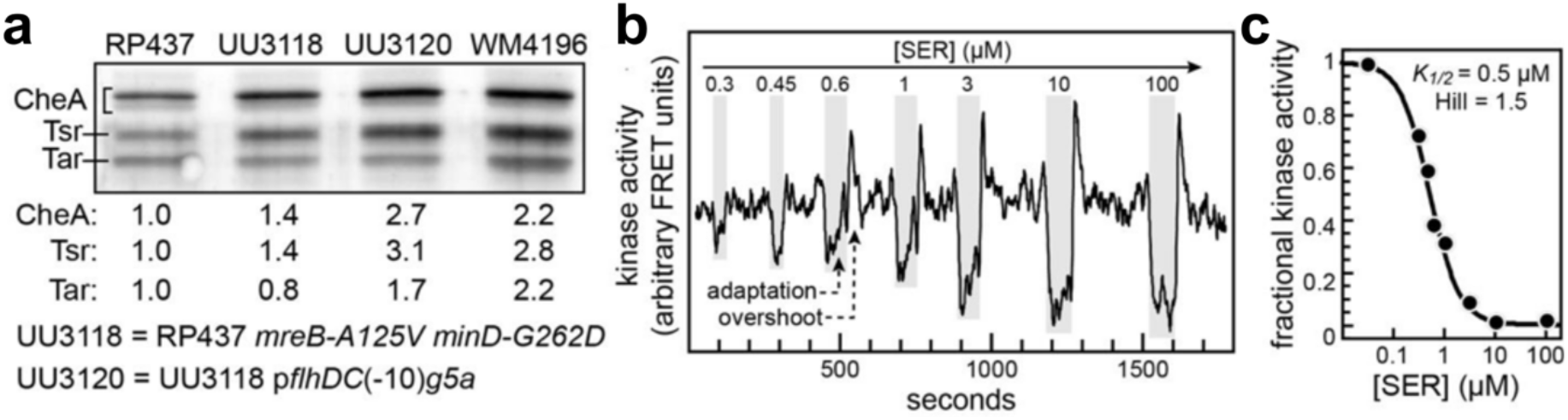
Chemotaxis proteins and signaling activities in WM4196 minicells. a) Tsr, Tar, and CheA levels in WM4196. Cell extracts were prepared and analyzed by SDS-PAGE and western blotting as described in Methods. Band intensities were determined by densitometry and normalized to the RP437 values. b) FRET analysis of serine signaling in WM4196 minicells. WM4196 minicells expressing the CheY-YFP/CheZ-CFP FRET reporter pair (see Methods) were immobilized on polylysine-coated coverslips and mounted in a microscope flow chamber.^27^ The horizontal trace follows the ratio of YFP to CFP emission counts, a measure of CheA kinase activity in the cells. Serine stimuli were applied to the cells during the intervals marked by gray rectangles. The magnitude of the drop in YFP/CFP value reflects the fraction of CheA activity inhibited by a serine stimulus. Maximal CheA activity in the cells is defined by the YFP/CFP drop in response to a saturating serine concentration (*e.g.*, 10 or 100 µM). These minicells contain the CheR and CheB adaptation enzymes which act to attenuate a serine-induced drop in kinase activity. A net methylation increase of the serine receptor during the adaptation phase produces a spike or overshoot in kinase activity upon serine removal that rapidly returns to the pre-stimulus kinase activity baseline.^26^ c) Hill fit of serine dose-response data. The fractional inhibition values from the experiment in panel (b) were fitted to a multi-site Hill function to determine the response *K*_*1/2*_, a measure of receptor sensitivity, and the Hill coefficient, a measure of response cooperativity.^44^

Third, to determine whether the arrays in WM4196 cells support chemotaxis, we examined WM4196 colony morphology on semi-solid agar motility plates. Despite apparent up-regulation of chemotaxis and, presumably, flagellar genes, we found that WM4196 performed quite poorly in soft agar chemotaxis tests compared to RP437 derivatives UU3118 and UU3120 (Supplementary Fig. 2). The observed migration difficulties of WM4196 could be due to the fact that receptor arrays reside mainly at the poles of cells. Because WM4196 mother cells efficiently bud minicells at their poles, the minicells probably contain most of the receptor arrays, leaving the mother cells with relatively few. However, since basal bodies form at random locations around the cell,^25^ few minicells are likely to have both a receptor array and a flagellar motor. Regardless, any minicells capable of chemotaxis would not contribute to colony expansion because they cannot reproduce. Perhaps the chemotaxis defects of their RP437 counterparts are less severe (Supplementary Fig. 2) because those strains bud minicells less efficiently and thereby have more array-containing mother cells during colony growth.

Finally, to ask whether the receptor arrays in WM4196 minicells are capable of detecting and responding to attractant stimuli, we employed a FRET assay (see Methods) that monitors receptor-coupled CheA kinase activity in intact cells.^26,27^ WM4196 minicells detected serine with high sensitivity (*K*_*1/2*_ = 0.4 µM) and moderate cooperativity (Hill coefficient = 2.7), values comparable to RP437 responses (Fig. 3c). The minicell responses exhibited hallmarks of sensory adaptation (response decay and activity overshoot; Fig. 3b) consistent with a normal complement of the CheR and CheB receptor-modifying enzymes. In summary, the receptor arrays in WM4196 minicells function comparably to those in RP437 cells. Therefore, we set out to calculate a 3D cryo-ET map of the CSU in WM4196 minicells.

In our cryo-ET reconstructions of individual minicells, the chemosensory array followed the curved surface of the inner membrane, providing the ensemble of views required for an isotropic 3D reconstruction by subtomogram averaging (see Methods, Supplementary Fig. 3, Supplementary Fig. 4). Subtomogram averaging in Dynamo^28–30^ resulted in a ~16 Å resolution 3D density map of a hexameric arrangement of three CSUs (Fig. 1c, Supplementary Fig. 3), from which we extract one for modelling and interpretation (Fig. 4, Supplementary Fig. 3). The most striking difference between our map and previously reported structures^4–8^ is the complete and continuous receptor density, showcasing the entire assembly from the periplasmic ligand-binding domain to CheA and CheW at the cytoplasmic signalling tip (Fig. 4a, Supplementary Fig. 3). Indeed, densities corresponding to the periplasmic domain, TM four-helix bundle, and HAMP domain were not resolved in previous array studies, which had been attributed to inherent flexibility of the CSU.^3^ Although our current map probably derives from a mixture of receptors with a range of adaptational modifications and signalling states, both the HAMP and MH bundles seem to be relatively static. Thus, our 3D reconstruction demonstrates that complete CSUs are amenable to visualisation by cryo-ET. The WM4196 minicell system, therefore, represents a valuable tool for elucidating structure-function relationships in CSUs, for instance through the imaging of arrays in different mutationally-imposed signalling states.

**Figure 4:**
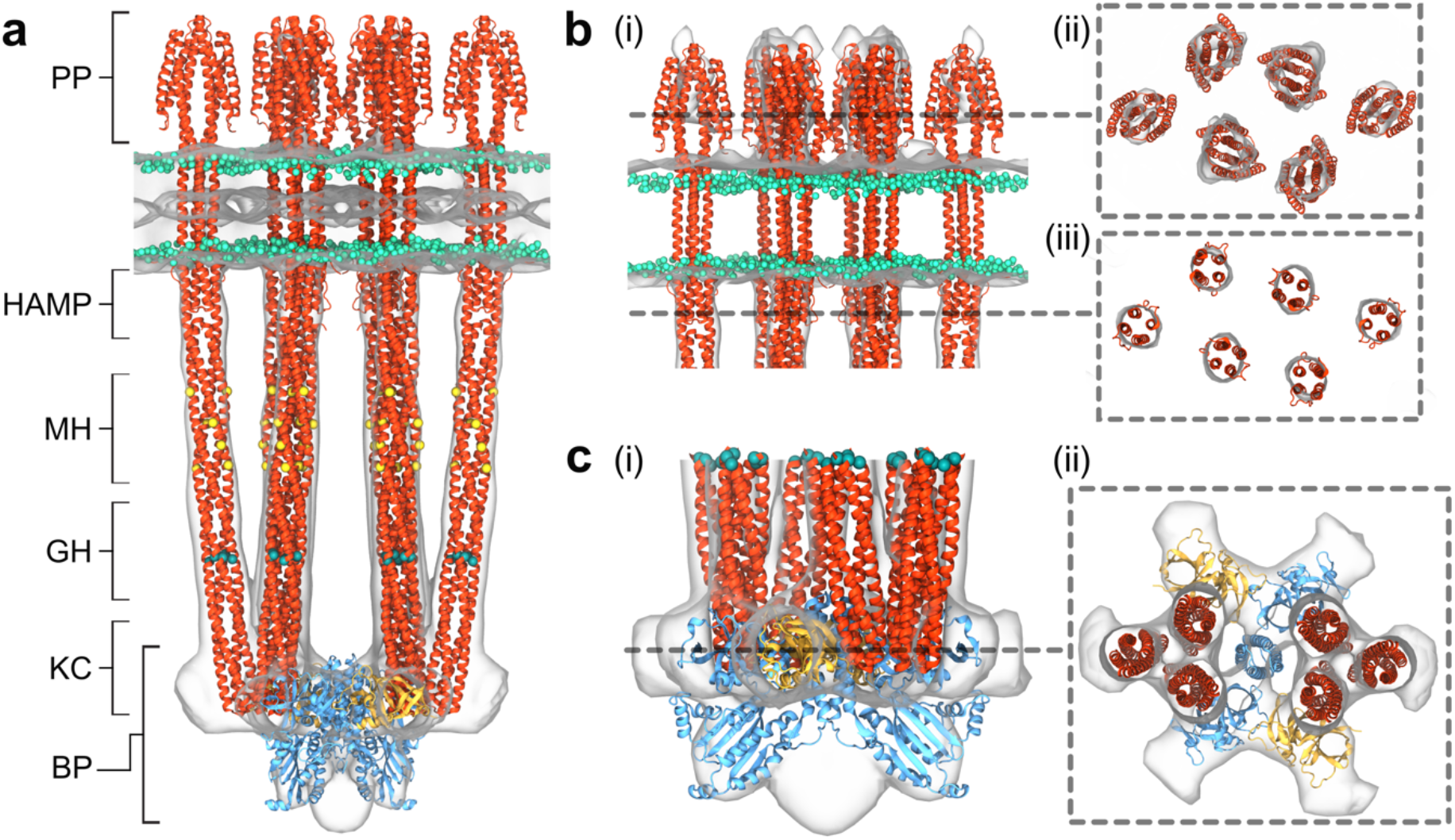
Atomistic model of complete *E. coli* core signalling unit. (a) Overlay between atomistic CSU model and 3D density map shown with surface representation. Receptors are depicted in red, CheA in blue, and CheW in gold. Membrane headgroups, methylation sites, and glycine hinge residues are shown as cyan, yellow, and teal spheres respectively. Regions corresponding to the periplasmic ligand-binding domains (PP), the HAMP domains, methylation-helix bundles (MH), flexible regions containing the glycine hinge (GH) and kinase control (KC) modules of the receptors are labelled, as well as the kinase baseplate (BP). (b) Zoom of membrane and membrane-proximal receptor domains from side (i) and at vertical slices taken through the periplasmic (ii) and HAMP domains (iii). To visualise the densities corresponding to the periplasmic domains, the map threshold is lowered in (i) and (ii). (c) Zoom of CheA/W baseplate region from side (i) and top (ii). In (i) the map and model have been rotated by 45^°^ to highlight excess density between CheA.P4 domains.

To enable spatial localisation of each signalling component and their individual domains in the 3D density map, we constructed an atomistic model of the complete *E. coli* CSU, using the density map to explicitly refine model tertiary structure (see Methods, Fig. 4a). The resulting molecular model provides several new structural insights throughout the CSU into MCP structure and organization. In particular, visualisation of the receptor ligand-binding domains allows us to describe for the first time the periplasmic organisation of the MCPs (Fig. 4b). Intriguingly, we observe in this region that receptors within a given ToD are situated as close or closer to those from neighboring ToDs than those in the same ToD (Fig. 4b, Supplementary Fig. 4), suggesting that minor diffusion within the membrane could give rise to ToD interactions both within and between CSUs.^31^ The impact of such interactions on signalling and cooperativity has yet to be systematically explored and should now be amenable to mutational and cross-linking analyses using WM4196-derived strains. Furthermore, previously documented kinks between the HAMP and the MH bundles as well as at the glycine hinge^32^ are clearly not required to avoid structural clashes between neighbouring MCPs (Fig. 4a), although gradual bending in these areas could play a role in transitions between signalling states.^8^ Overall, the symmetry axes of the ToDs are separated by 7.4 nm, suggesting an array lattice constant of ~12.8 nm which is consistent with inter-particle distances in the tomograms.(Supplementary Fig. 5).

In the present model, the CSU baseplate, which comprises densities corresponding to the MCP signalling tip, the CheA kinase dimer, and the CheW coupling protein (Fig. 1b), is globally consistent with published cryo-ET-based models.^4–6^ The tips of the individual receptor dimers embedded in the CheA.P5-CheW lattice are well defined, and the known MCP-CheA.P5, MCP-CheW, and CheA.P5-CheW interfaces critical for assembly are preserved (Fig. 4c). The CheA.P3 dimer sits between and parallel to the two ToDs^4,6^, while the CheA.P4 ATP-binding domain is localized underneath CheA.P5 (Fig. 4c) as seen in previous structures.^4,8,33^ Intriguingly, a substantial amount of density situated below and between the two P4 domains remains unaccounted for (Fig. 4c). Several lines of evidence suggest that the P1 and P2 domains of CheA undergo dynamic changes during signalling, and it has been proposed that these domains are sequestered in the kinase-OFF state.^7,34^ In addition, NMR experiments have mapped a non-productive P1-P4 binding mode to this region of P4 in *T. maritima* CheA.^35^ Therefore, the unknown density may correspond to either one or both of the CheA.P1 domains and could also include CheA.P2. In general, the role of dynamics in CheA signalling and the mechanisms of receptor regulation are matters of controversy. Thus, our WM4196 minicell system provides a much-needed tool for assessing these issues experimentally. Finally, we note that while we focus here on a single CSU, our approach may be extended to probe long-range and cooperative signal transduction between CSUs, for example by analysing array packing in different signalling states. Thus the WM4196 strain also offers a novel tool for investigating the molecular mechanisms underlying the remarkable signal amplification and wide dynamic range of the chemosensory system.^9,36^

Thanks to their small size and structural integrity, the WM4196 minicells presented in this work reveal themselves as suitable for cryo-ET studies in general. They are also particularly appropriate for higher resolution analysis of chemosensory arrays due to their higher expression level of chemosensory components. Moreover, because WM4196 minicells bud closer to the bacterial pole, they contain a higher percentage of arrays than larger minicell strains, enabling more efficient acquisition of usable tilt-series for subsequent subtomogram averaging. Here we have shown that properties of the WM4196 strain lead to a significant improvement in the resolution of the *in situ* array structure and an unparalleled definition of the MCP structure, including the periplasmic ligand binding, HAMP, and the MH bundle domains that play a vital role in converting ligand-induced conformational rearrangements into CheA control. The results obtained in the present work can now be fruitfully combined with higher resolution studies of *in vitro* reconstituted chemosensory arrays^4^ and compared with cryo-ET reconstructions from other bacteria such as *Vibrio cholerae*, *Rhodobacter sphaeroides* and *Caulobacter crescentus*.^37–40^ Finally, these small and healthy minicells open up new opportunities for the study of *E. coli* macromolecular complexes located near the cell poles.

## Methods

### Design and genetic characterisation of the WM4196 strain

Strain WM4196 is a distant derivative of a minicell mutant first described over 50 years ago.^20^ This prototype strain, containing the *minD* allele, a *dnaB* ts allele, as well as several auxotrophic markers, was converted to *dnaB*+ and prototrophy by conjugation with a prototrophic Hfr donor strain.^22^ The F’ was then cured to make strain x1411, which was obtained from the *E. coli* Genetic Stock Center, Yale University (CGSC#6397). The *mreB*-A125V allele^21^ was further subsequently introduced into x1411 by cotransduction with a tightly linked *yhdE::cat* chromosomal allele to create the WM4196 strain.

### DNA sequencing and strain constructions

WM4196 chromosomal DNA was PCR-amplified and sequenced with primer pairs specific for the *minCDE*, *flgM*, and *flhDC* loci. Strain UU3118 (RP437 *mreB-A125V minD-G262D*) was constructed by (a) introducing the *mreB* allele from WM4196 into RP437 by phage P1-mediated cotransduction with the linked *∆yhdE::cat* allele; and (b) introducing the *minD-G262D* mutation with homologous recombination-mediated insertion and replacement steps.^41^ Strain UU3120 is a derivative of UU3118 that carries the *pflhDC(−10)g5a* allele of WM4196 introduced by two-step insertion/replacement.

### Measuring cellular levels of chemotaxis proteins

Plasmid pRR48^42^ was introduced into RP437, WM4196, and UU3111 to express the β-lactamase (Bla) protein (an internal reference for standardizing the relative levels of other cell proteins). Cells were grown at 37°C in L broth (10 g/L tryptone, 5 g/L yeast extract, 5 g/L NaCl) containing 25 µg/ml ampicillin and harvested at mid-exponential phase (OD_600_ = 0.5−0.6). Approximately equal numbers of cells (based on culture OD) were used to prepare lysates. Cells were pelleted by centrifugation, washed once with motility buffer (10 mM K-PO4 [pH 7.0], 0.1 mM K-EDTA) and lysed in SDS-PAGE sample.^43^ Samples were analyzed in 11% polyacrylamide-SDS gels and MCP, CheA and Bla bands were detected by immunoblotting with a mix of three polyclonal rabbit antisera. Bands were visualized with a Cy5-labeled goat anti-rabbit antibody and quantified with a fluorescence imager.

### In vivo FRET kinase assay

This method is described in detail in previous publications.^27,44^ For the present study, we introduced into WM4196 plasmid pVS88^26^, which expresses fusion proteins CheZ-CFP (FRET donor) and CheY-YFP (FRET acceptor) under IPTG-inducible control. The plasmid-carrying strain was grown at 37°C to OD_600_=0.65−0.7 in L broth (see above) containing 25 µg/ml ampicillin and 1 mM IPTG. (The high IPTG concentration compensates for elevated levels of untagged CheY and CheZ proteins expressed from the WM4196 chromosomal genes.) 300 ml of cell culture were centrifuged at ~8600 X g for 25’ at 4°C. All subsequent centrifugation steps were also carried out at 4°C. The supernatant was transferred to fresh tubes and centrifuged at ~14,500 X g for 25’. The sample pellets were pooled and resuspended in ~1 ml of motility buffer (see above) and carried through a second round of differential centrifugation (9000 X g for 10’; 21000 X g for 20’). The final minicell pellet was resuspended in ~100 µl motility buffer and applied to a polylysine-coated cover slip for the FRET assay.

### WM4196 minicell purification for cryo-ET imaging

WM4196 minicells were grown in L broth supplemented with 34 µg mL^−1^ chloramphenicol for 12 hours. This culture was used to inoculate larger L broth cultures (without antibiotics) to an OD_600_ value of 0.075. Bacteria were grown at 37 °C for 4h (the culture was grown to an OD_600_ value of 0.5 before being left to grow for a further 2h. Final OD_600_ value was 1.75). The culture was centrifuged at 8683 g for 20 minutes at 4 °C, the supernatant carefully transferred to another centrifuge tube and centrifuged again at 8683 g for 20 minutes at 4 °C. The resulting supernatant was transferred to another centrifuge tube and centrifuged at 41500 g for 20 minutes at 4 °C. This time, the supernatant was discarded and the pellet gently resuspended in the residual supernatant from the centrifuge tube. This suspension was then centrifuged at 5500 g for 5 minutes at 4 °C, and the supernatant transferred to another centrifuge tube to be centrifuged at 16900 g for 15 minutes at 4 °C. The resulting minicell pellet was resuspended in LB and the minicell suspension was placed at 4 °C.

### Cryo-electron microscopy imaging and cryo-ET data acquisition

Minicell solution was supplemented with 10 nm colloidal protein A-gold particles (Cell Microscopy Core, University Medical Center, Utrecht, The Netherlands). R 2/1 or R 3.5/1 on 300 mesh Cu/Rh grids (QUANTIFOIL) were glow discharged for 45 s at 25 mA. 3 μL of the sample was applied to grids and plunge frozen in liquid ethane using a Vitrobot Mark IV (Thermo-Fisher Scientific). Grids were stored under liquid nitrogen until imaging. Screening of the grids was performed on an FEI F20 microscope at 200 keV. Tilt-series acquisition was performed on a Titan Krios transmission electron microscope operated at 300 keV, through a Gatan Quantum energy filter and Volta phase plate onto a K2 Summit direct electron detector, using SerialEM software.^45^ For both search and navigational purposes, low-magnification montages were acquired. For tilt-series acquisition, a magnification which produced a calibrated pixel size of 2.24 Å was selected. Data were collected at tilt-angles between −60° and +60° in 2° increments in a grouped dose-symmetric tilt scheme (0, +2, +4, +6, +8, +10, −2, −4, −6, −8, −10, 12, 14, 16, 18, 20, −12, −14, −16, −18, −20 etc).^46^ Images at each tilt-angle were acquired as movies comprising 5 frames. Target total dose for tilt-series acquisition was 60 e^−^ Å^−2^s^−1^.

### Image Processing

#### Pre-Processing

Frames of movies corresponding to each tilt-angle were aligned and motion-corrected using MotionCor2^47^ to mitigate the deleterious effects of beam-induced sample motion. Tilt-series were assembled using the newstack command from IMOD.^48^

#### Tomographic Reconstruction

Tilt-series were aligned, binned by two and tomograms were reconstructed by weighted back-projection using the IMOD software package.^48^ Six tomograms of WM4196 minicells were selected for further processing.

#### Initial Reference Generation

Particles were identified and picked using the Dynamo software package.^28–30^ Sub-volumes with a side length of 573 Å were extracted and initial orientations for each particle were estimated by modelling a surface following the curvature of the chemosensory array in the tomogram and imparting an orientation onto each particle which corresponded to the normal to the modelled surface at the point closest to the particle. These orientations were further refined manually, and these coarsely-oriented particles were then averaged to produce an initial reference. The particles were subsequently locally aligned, constraining the angular search to a 60° cone around the initial estimate for the orientation, and averaged to produce initial references.

#### Particle Picking

The following analysis was performed in Dynamo.^29^ Surfaces were modelled following the curvature of the chemoreceptor arrays visible inside the minicells and a set of initial positions and orientations was generated from this surface with an average distance of 30 Å between each position. Sub-volumes with a side length of 573 Å were extracted at each of these positions and each was aligned to the initial reference with an allowed translational freedom of 60 Å in each of the x-, y- and z-dimensions. Analysis of the post-alignment positions revealed an ordered hexagonal array. Duplicate particles (defined as particles within 10 Å of each other) were collapsed into one final position. The particle positions were then cleaned by selecting only those which had greater than three nearest-neighbours at the expected distance of 120±20 Å.

#### Subtomogram Averaging

Alignment and averaging of the particles was performed in the Dynamo software package.^28^ An iterative global alignment of the particle positions and orientations was performed starting from the initial reference already generated, for which alignments were performed inside a mask encompassing multiple core-signalling units in the chemosensory array and the inner membrane of the WM4196(DE3) minicell. These positions and orientations were then further locally refined inside a mask containing only three core-signalling units, without the membrane^10^, to produce a final reconstruction.

#### Post Processing

Two separate half-maps were generated from groups of particles coming from different tomograms to allow estimation of the resolution of the final reconstruction. The fourier shell correlation of these maps inside a mask, containing three core-signalling units and no membrane, drops below 0.143 at a spatial frequency corresponding to a resolution of 16 Å. The maps were subsequently subject to localised resolution estimation in RELION^49^ with a sampling rate of 20 Å and locally filtered to the estimated resolution of the map. The local-resolution filtered map was aligned to a C2-symmetric reference centered on one core-signalling unit and then itself symmetrised around the C2-axis to give a map centered on one core-signalling unit. Masks for FSC calculation and map visualisation were calculated using a combination of Chimera^50^, Dynamo^28^ and RELION.^49^

#### All-atom model generation

Coordinates for *E. coli* CheA.P3.P4.P5 and CheW were derived from existing *T. maritima* crystal structures^33,51^ using template-based homology modeling, while a complete model of the *E. coli* Tsr was assembled from existing crystal structures of the ligand binding^52^ and cytoplasmic domains^53^ as well as a HAMP homology model based on a crystal structure of *A. fulgidus* HAMP.^54^ The TM four-helix bundle was modelled using existing cross-linking data between the individual TM helices^55,56^ as well as constraints implied by the ligand binding and HAMP domain structures, the full details of which are the subject of an upcoming publication. The complete Tsr model was embedded in an atomistic lipid bilayer and subjected to an extensive structural equilibration using molecular dynamics simulation. The component models were then assembled via rigid docking into the density map and refined using Molecular Dynamics Flexible Fitting^57,58^ as described previously.^4,59^

#### Data availability

The Cryo-ET derived map has been submitted to the EMDB with accession code EMD-10160.

## Acknowledgements

This work has received funding from a European Union’s Horizon 2020 research and innovation programme under grant agreement No 647784 to IG. Research in the JSP and WM labs was supported by grants GM19559 and GM131705, respectively, from the U.S. National Institute of General Medical Sciences. This work was also supported by the U.S. National Institutes of Health grant P41GM104601 to CKC and ZL-S, the U.S. National Science Foundation grant PHY1430124 to CKC and ZL-S, and the U.K. Biotechnology and Biological Sciences Research Council grant BB/S003339/1 to CKC and PJS. We acknowledge Diamond Light Source for access and support of the cryo-EM facilities at the UK’s national Electron Bio-imaging Centre (eBIC), funded by the Wellcome Trust, MRC and BBRSC. Cryo-ET data acquisition has been supported by iNEXT, grant number 653706 (PID:2626 to IG), funded by the EU Horizon 2020 programme. For initial minicell characterisation and grid screening, we used the platforms of the Grenoble Instruct-ERIC Center (ISBG: UMS 3518 CNRS-CEA-UGA-EMBL) with support from FRISBI (ANR-10-INSB-05-02) and GRAL, a project of the University Grenoble Alpes graduate school (Ecoles Universitaires de Recherche) CBH-EUR-GS (ANR-17-EURE-0003). IBS acknowledges integration into the Interdisciplinary Research Institute of Grenoble (IRIG, CEA). The IBS electron microscope facility is supported by the Rhône-Alpes Region, the Fondation pour la Recherche Médicale (FRM), the fonds FEDER, the Centre National de la Recherche Scientifique (CNRS), the Commissariat à l’Energie Atomique et aux Energies Alternatives (CEA), the University of Grenoble Alpes, EMBL, and the GIS-Infrastructures en Biologie Santé et Agronomie (IBISA). Molecular dynamics simulations were performed on the Blue Waters supercomputer, which is supported by the National Science Foundation (OCI-0725070 and ACI-1238993) and the state of Illinois as part of the Petascale Computational Resource Grant (ACI-1713784). We are particularly grateful to Daniel Clare, Alistair Siebert and Andrew Howe for help with data acquisition at eBIC, and to Daniel Castaño-Diez for help and discussions on image processing and for development of Dynamo. We thank Guy Schoehn for establishing and managing the IBS Grenoble cryo-electron microscopy platform and for providing training and support. We are grateful to Aymeric Peuch for help with the usage of the joint IBS/EMBL Grenoble EM computing cluster.

## Author Contributions

IG designed and supervised the study. WM created the WM4196 minicell strain. AB, MB and KH purified the minicells, MBV, AB and IG screened them by cryo-EM. PA, WM and JSP created the RP437 derivatives, and characterized them and the WM4196 strain. AB, AD and IG collected cryo-ET data, AB analysed cryo-ET data with input from AD and IG. CKC constructed and refined the atomistic model in discussion with ZL-S and PJS and interpreted it in the context of the cryo-ET density with input from AB and IG. AB, CKC, PA, JSP and WM prepared the figures and movies. IG wrote the manuscript with significant input from AB, CKC, PJS, WM and JSP and contribution of all authors.

