## Supplementry Information for "Complete structure of the core signalling unit of the *E. coli* chemosensory array in an optimised minicell strain"

**Supplementary Figures 1 to 5**

**Supplementary Movie 1**

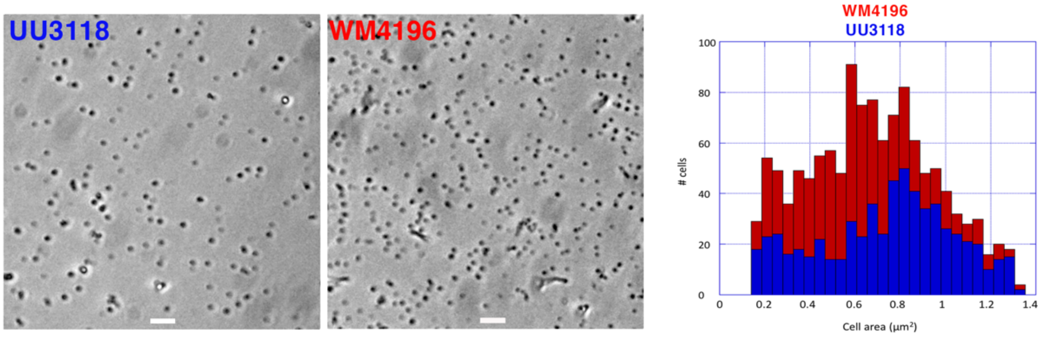


**Supplementary Figure 1: Size comparison of WM4196 and UU3118 minicells.** Minicells from WM4196 and UU3118 cultures were purified by differential centrifugation as described in Methods for the FRET kinase assay, immobilized on a polylysine-coated coverslip, and imaged at the same magnification by phase contrast microscopy (top panels, scale bar = 4 µm). After passing the images through a threshold filter to facilitate cell measurements, ImageJ software and its plug-in MicrobeJ[1] were used to identify hundreds of minicells, define their edges and calculate their 2-D surface areas from their dimensions. To minimize the background of non-minicells or inaccurate measurements, we implemented arbitrary cutoffs of areas below 0.15 $\mu m^2$ and above 1.35 $\mu m^2$. The measured minicell areas of each strain were then plotted as frequency distributions (bottom panel) using Kaleidagraph (Synergy Software). The measured areas may be overestimated due to optical artifacts and resolution limits but should be comparable between strains.

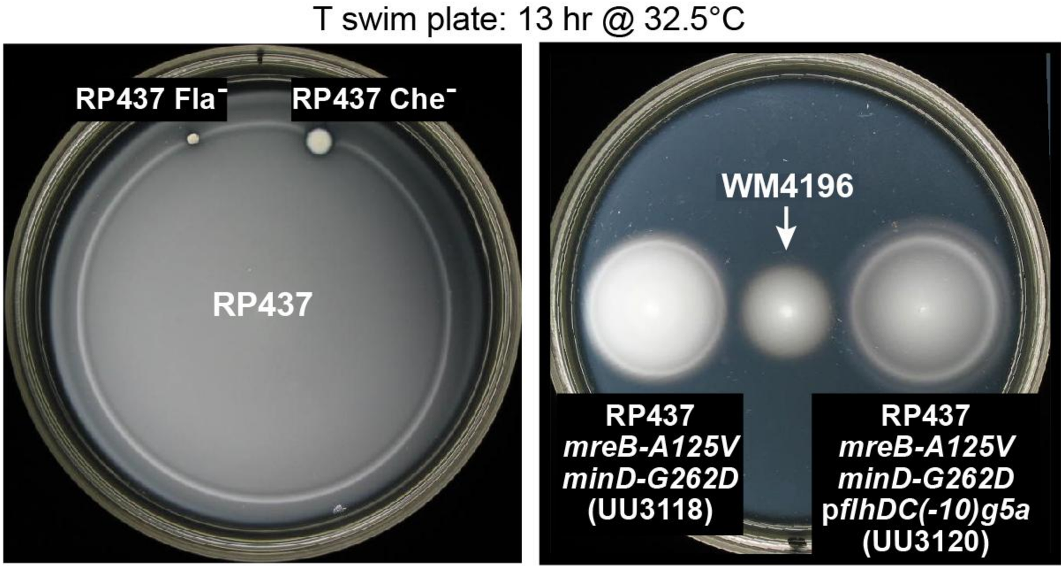


**Supplementary Figure 2: Chemotaxis performance of RP437 and WM4196 strains.**

Cells were streaked on T hard plates (T broth plus 14 g/L agar) and grown for 24 hours (WM4196) or 18 hours (RP437 strains) at 37°C, Colonies were picked to T swim plates (T broth plus 2.5 g/L agar) and incubated at 32.5°C for 13 hours. Chemotactic colonies exhibit one or more dense bands or rings of cells that track metabolism generated nutrient gradients in the plate.

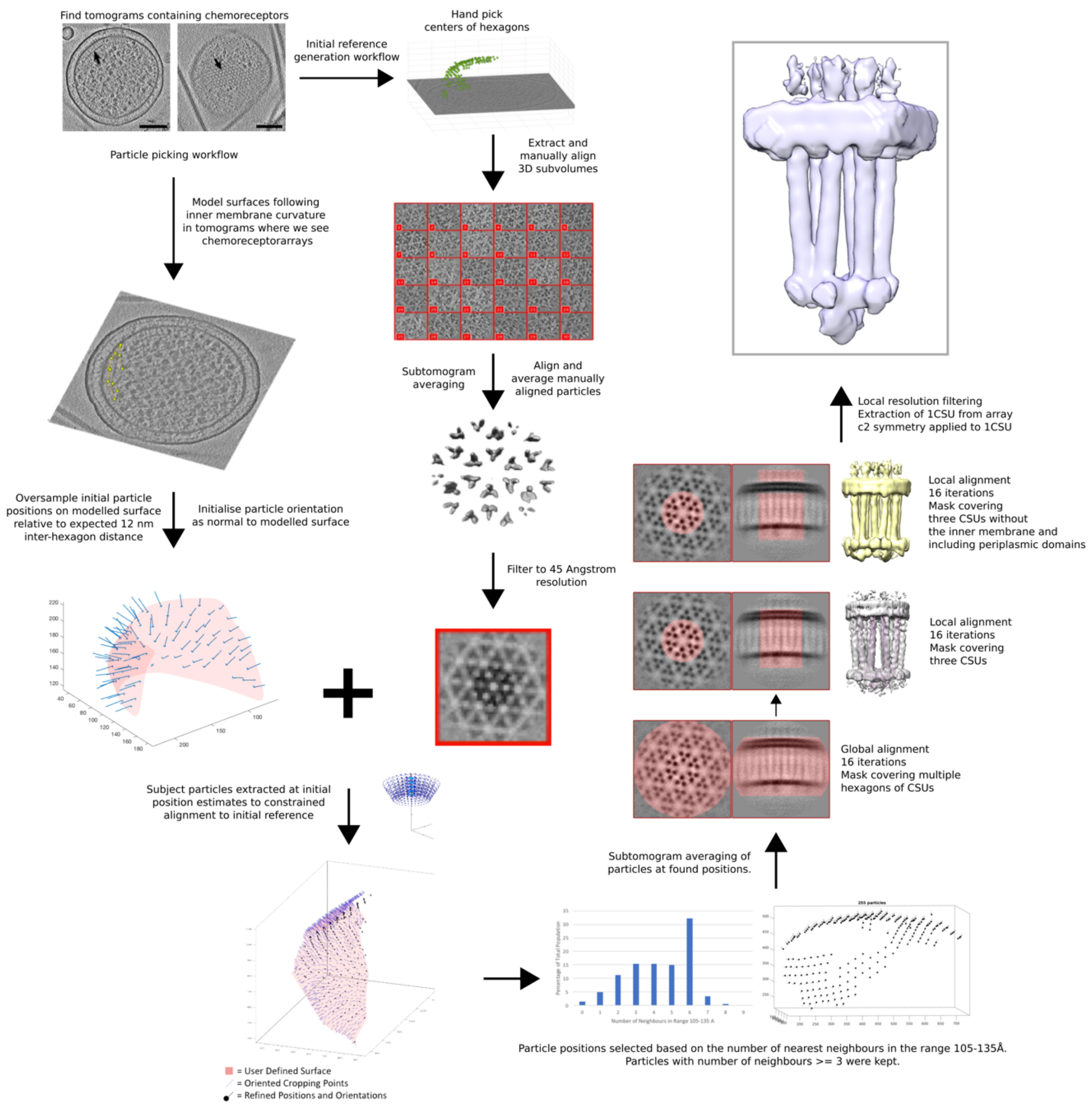


**Supplementary Figure 3: Flowchart of the cryo-ET image processing pipeline.** A generalized overview of the subtomogram averaging workflow used to obtain the *in situ* reconstruction of a complete CSU presented in this manuscript using Dynamo.[2,3] For a full description of the image processing workflow, see 'Methods'.

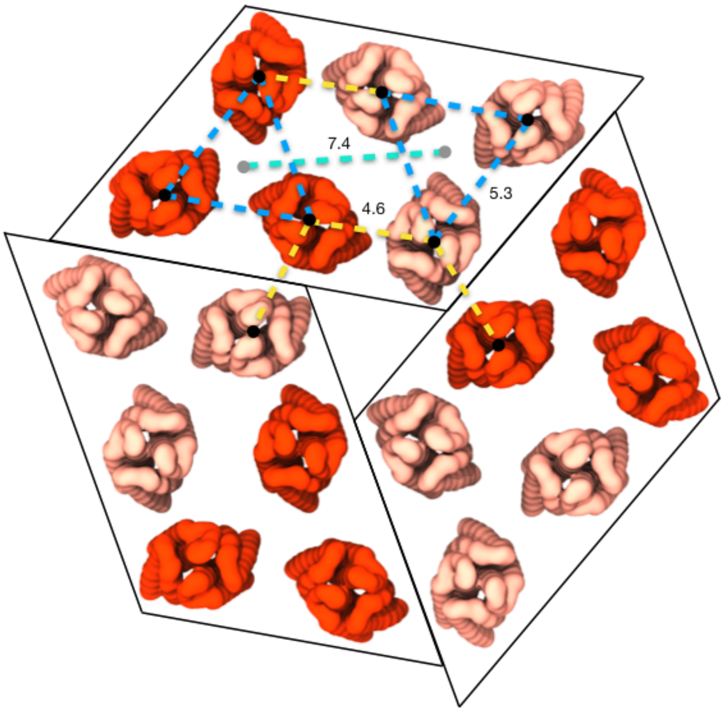


**Supplementary Figure 4: Extended organization of receptor periplasmic domains**

Distances between receptor periplasmic domains within and between core signalling units (CSUs) in the extended array. Individual core signalling units are boxed by a parallelogram with comprising receptor trimers of dimers (ToDs) colored in red and salmon. Distances are given in nanometers and are computed between the centers-of-mass of the receptor periplasmic domains (black circles) and the ToD symmetry axes (gray circles). Dashed lines of the same color denote equal distances. The model of the extended array architecture shown here was constructed by rigidly docking the MDFF-refined CSU model (Fig. 4) into a the density map for the array centered on a hexagonal arrangement of 3 CSUs.

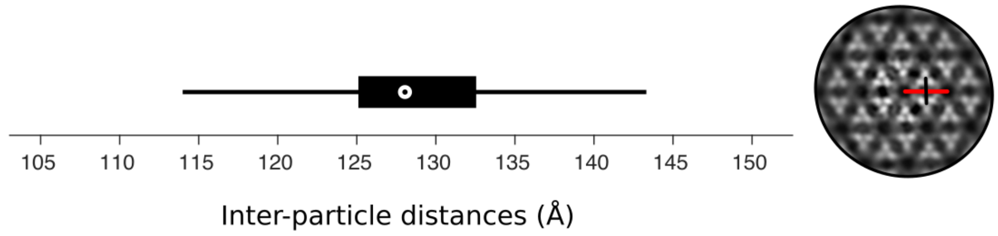


**Supplementary Figure 5: Distribution of inter-particle distances in tomograms**

A box plot to show the distribution of inter-particle distances in tomograms of WM4196 minicells. The central mark indicates the median (128 Å), the left and right edges of the box indicate the 25$^{th}$ and 75$^{th}$ percentile respectively. Whiskers extend to approximately 2.7σ. Particle positions (CheA/W ring centers) were found by subtomogram averaging. Inter-particle distances measured correspond to the red line shown on a 5 nm projection through the density of our reconstruction. The 7.4 nm inter-ToD distance corresponds to the black line perpendicular and above the red line.

**Supplementary Movie 1: WM4196 tilt-series, tomogram and the derived map and model.**

A movie to show a tilt-series which contributed to the final reconstruction, the corresponding tomographic reconstruction and the in situ core-signalling unit density, the derived MDFF equilibrated model and the fit of the model in the density. The threshold for the surface representation of the model is lowered during the rotation to accentuate the presence of the periplasmic ligand-binding domains.
